# NBR1 directly promotes the formation of p62 – ubiquitin condensates via its PB1 and UBA domains

**DOI:** 10.1101/2020.09.18.303552

**Authors:** Adriana Savova, Julia Romanov, Sascha Martens

## Abstract

Selective autophagy removes harmful intracellular structures such as ubiquitinated, aggregated proteins ensuring cellular homeostasis. This is achieved by the encapsulation of this cargo material within autophagosomes. The cargo receptor p62/SQSTM1 mediates the phase separation of ubiquitinated proteins into condensates, which subsequently become targets for the autophagy machinery. NBR1, another cargo receptor, is a crucial regulator of condensate formation. The mechanisms of the interplay between p62 and NBR1 are not well understood. Employing a fully reconstituted system we show that two domains of NBR1, the PB1 domain which binds to p62 and the UBA domain which binds to ubiquitin, are required to promote p62-ubiquitin condensate formation. In cells, acute depletion of endogenous NBR1 reduces formation of p62 condensates, a phenotype that can be rescued by re-expression of wild-type NBR1, but not PB1 or UBA domain mutants. Our results provide mechanistic insights into the role of NBR1 in selective autophagy.

## Introduction

Macroautophagy (hereafter autophagy) is a conserved intracellular process, which aids cellular homeostasis by the disposal of harmful structures such as protein aggregates, damaged organelles and intracellular pathogens (Noboru Mizushima & Komatsu, 2011; Randow, MacMicking, & James, 2013; Zaffagnini & Martens, 2016). Defects in autophagy have been linked to a plethora of diseases including cancer and neurodegeneration (Levine & Kroemer, 2019). The degradation of the harmful material, referred to as cargo, is mediated by its encapsulation within *de novo* formed double membrane vesicles, termed autophagosomes, which fuse with lysosomes wherein the cargo is degraded. Autophagosome formation is mediated by the autophagy machinery (N Mizushima, Yoshimura, & Ohsumi, 2017; T. Nishimura & Tooze, 2020). In selective autophagy, during which cargo is specifically targeted, this machinery is recruited to the cargo by cargo receptors such as p62/SQSTM1, NBR1, NDP52 and optineurin (Ravenhill et al., 2019; Smith et al., 2018; Turco, Fracchiolla, & Martens, 2020; Turco et al., 2019; Vargas et al., 2019; Yamano et al., 2020). Many of the various cargo receptors in mammalian cells, including p62 and NBR1, recognize the cargo via its ubiquitination (Vladimir Kirkin & Rogov, 2019).

A major function of p62 is the degradation of ubiquitinated, misfolded proteins by selective autophagy. In this process it plays several roles (Danieli & Martens, 2018) (Johansen & Lamark, 2020; Sánchez-Martín, Saito, & Komatsu, 2019). First, it mediates the phase separation of ubiquitinated proteins into larger condensates (Bjørkøy et al., 2005; Sun, Wu, Zheng, Li, & Yu, 2018; Zaffagnini et al., 2018). Subsequently, it contributes to the local formation of autophagosomes around these condensates by recruiting the scaffolding FIP200 protein (Turco et al., 2019). Finally, it links the cargo to the nascent autophagosomal membrane via its interaction with LC3 and GABARAP proteins, which decorate the forming autophagosomal membrane (Ichimura et al., 2008; Pankiv et al., 2007). p62 oligomerizes into filaments through its N-terminal PB1 domain (Ciuffa et al., 2015; Jakobi et al., 2020; Lamark et al., 2003). This oligomerization is required for its ability to phase-separate ubiquitinated proteins but also to avidly bind the LC3/GABARAP decorated autophagosomal membrane via its LC3 interacting region (LIR) motif and the ubiquitinated cargo via its C-terminal UBA domain (Sun et al., 2018; Wurzer et al., 2015; Zaffagnini et al., 2018).

In these processes, p62 is aided by the cargo receptor NBR1 (Fig 1A). Similar to p62, NBR1 binds LC3/GABARAP proteins via LIR motifs and ubiquitin via its UBA domain (Pankiv et al., 2007). NBR1 and p62 directly interact through their N-terminal PB1 domains (Jakobi et al., 2020; Lamark et al., 2003; Zaffagnini et al., 2018). In cells NBR1 colocalizes with p62 and its depletion results in fewer p62 condensates (V Kirkin et al., 2009; Sánchez-Martín et al., 2020). *In vitro*, NBR1 directly enhances the formation of p62-ubiquitin condensates (Zaffagnini et al., 2018). The specific mechanisms through which NBR1 promotes condensate formation and thus cargo degradation remain unclear.

**Figure 1.**
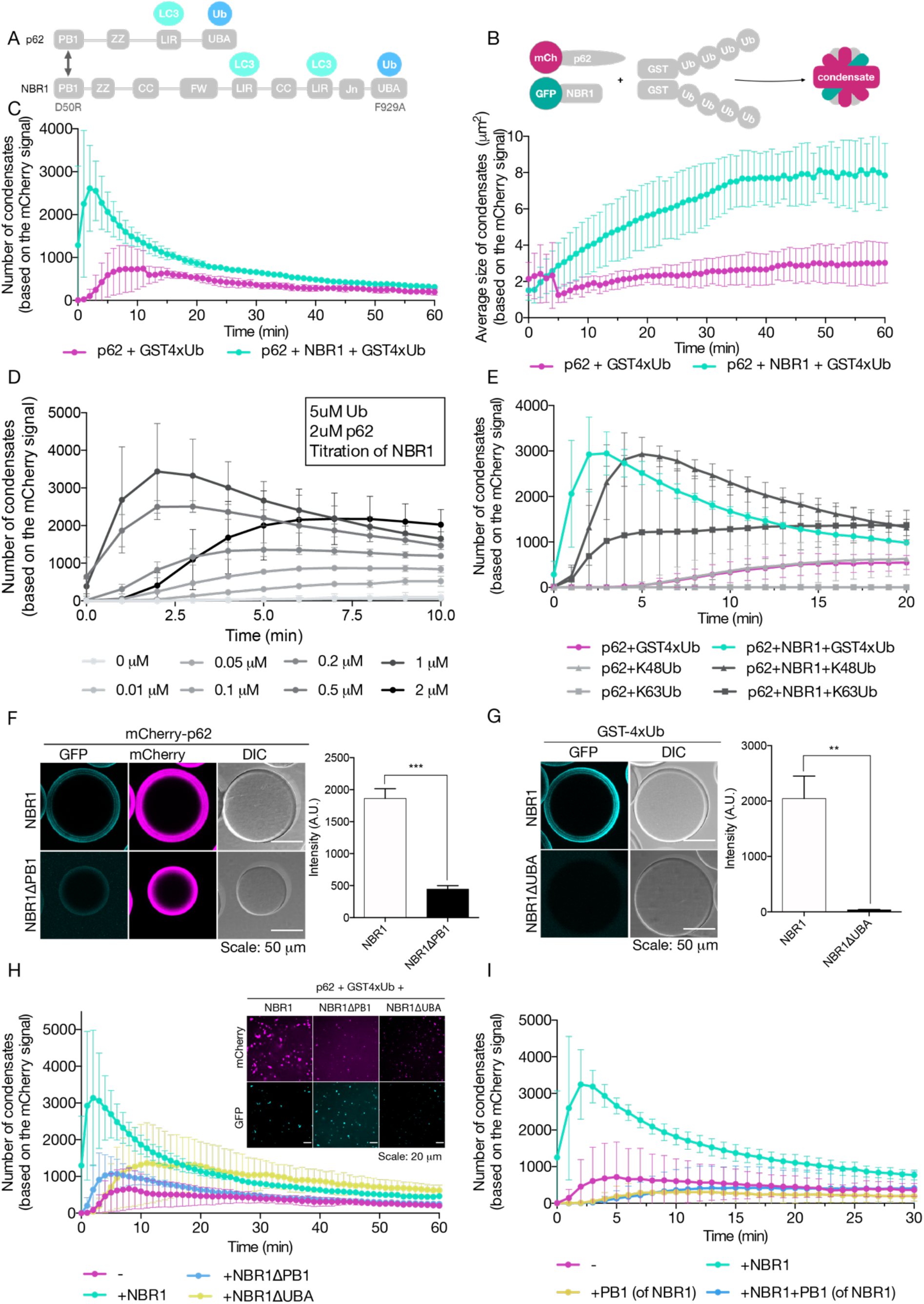
NBR1 regulates p62-cargo phase separation *in vitro*. **(A)** Schematic of the domain organization of the autophagy cargo receptors p62/SQSTM1 and NBR1. **(B)** Schematic of *in vitro* phase separation assays: mCherry-p62 is premixed with GFP-NBR1 in SEC buffer (25mM Hepes pH 7.5, 150mM NaCl, 1mM DTT). GST4xUb is added last to trigger condensation. **(C)** Quantification of a phase separation assay with 2 μM mCherry-p62, 2 μM GFP-NBR1 and 5 μM GST4xUb. The quantification is done based on the mCherry signal. Plotted graphs show total number of condensates per imaging field and average sizes of condensates. A quantification based on GFP, representative images and protein loading controls are available in Fig S1A, B. **(D)** Titration of NBR1 in the range of 0-2 μM into a condensate formation assay. All reactions contain 2 μM mCh-p62 and 5 μM GST4xUb. A control for equal protein loading can be found in Fig S1C. **(E)** Quantification of a phase separation assay with 5 μM GST4xUB or 5 μM *in vitro* synthesized K48-linked or K63-linked ubiquitin chains of variable length. Assay performed in SEC buffer with 2% BSA to simulate crowding conditions. The chains used in the reaction are loaded on a control Coomassie gel in Fig S1D. **(F, G)** Quantification of recruitment of 2uM GFP-NBR1 variants to RFP-trap beads saturated with mCherry-p62 **(F)** or to GST-trap beads saturated with GST-4xUb **(G). (H)** Quantification of a phase separation assay with 2 μM mCherry-p62, 2 μM GFP-NBR1 and 5 μM GST4xUb. Tested either full length NBR1 or ΔPB1 or ΔUBA variants. The quantification is based on the mCherry signal. Representative images of the reaction at timepoint 60 minutes are included. A silver gel controlling for protein purity is available in Fig S1E and a Coomassie gel showing the proteins loaded in the reaction in Fig S1F. **(I)** Phase separation assay with 2 μM mCh-p62 and 5 μM GST4xUb. In addition, 2 μM GFP-NBR1, 2 μM of the PB1 domain of NBR1 (PB1-NBR1) or both were tested on top and plotted. Equal protein loading is shown on a Coomassie gel in Fig S1G.

Here we employ a fully reconstituted assay, as well as cells containing endogenously tagged proteins coupled to acute auxin-induced degradation of NBR1, to study how NBR1 promotes the formation of p62-ubiquitin condensates. We show that *in vitro* NBR1 promotes the condensation of substrates modified with M1-(linear), K48- and K63-linked ubiquitin chains in a concentration dependent manner, with the optimal NBR1 concentration being substoichiometric to p62. By using PB1 and UBA domain mutants of NBR1, we show that NBR1 promotes p62-ubiquitin condensate formation via its PB1 domain mediated binding to p62. This interaction additionally equips the p62-NBR1 heterooligomeric complex with a high-affinity UBA domain provided by NBR1, allowing for more efficient cargo recognition.

## Results

### NBR1 modulates p62 - ubiquitin condensate formation *in vitro*

Previous work has shown the ability of p62 and ubiquitinated proteins to phase separate *in vitro* (Sun et al., 2018; Zaffagnini et al., 2018). It was further shown that NBR1 directly promotes phase separation of p62 with linear ubiquitin chains in a reconstituted system (Zaffagnini et al., 2018). How NBR1 promotes phase separation and if it also promotes phase separation of the more abundant and likely more physiologically relevant substrates modified with K48- and K63-linked ubiquitin chains is unknown. To answer these questions, we employed our *in vitro* phase separation assay (Fig 1B)(Zaffagnini et al., 2018). Consistent with our previous results, we observed that NBR1 alone did not induce phase separation but that it markedly enhanced the phase separation of p62 with GST fused to four M1-linked ubiquitin moieties (GST-4xUb), as judged from the number (left panel) and size (right panel) of the condensates (Fig 1C, Fig S1A). NBR1 was efficiently recruited to p62-containing condensates (Fig S1A). In order to test if there was an optimal molar ratio of NBR1 to p62 with respect to its phase separation promoting activity, we titrated NBR1 into phase separation assays containing p62 and GST-4xUb (Fig 1D). We observed that the promoting activity of NBR1 gradually increased with increasing concentrations, but that it dropped at an equimolar ratio. This suggests that a substoichiometric concentration of NBR1 relative to p62 is ideal for the promotion of condensate formation.

Next, we asked if NBR1 would also promote the phase separation of K48- and K63-linked ubiquitin chains (Fig 1E, Fig S1D). NBR1 promoted the phase separation of these chain types as well. There are differences in the degree to which the phase separation of the three substrates was promoted, but due to the different length of the chains (Fig S1D), we cannot distinguish between chain type and chain length. We conclude that NBR1 can enhance the phase separation of a broad spectrum of substrates by p62, which is consistent with a low degree of ubiquitin linkage specificity of its UBA domain (Walinda et al., 2014).

We went on to dissect which properties of NBR1 are required to promote phase separation. To this end, we expressed and purified two deletion mutants of NBR1 (Fig S1E). The first mutant lacked the PB1 domain and as expected from previous studies (Jakobi et al., 2020; Lamark et al., 2003) showed a severely reduced binding of NBR1 to p62 (Fig 1F). We also deleted the C-terminal UBA domain which was previously shown to mediate binding to ubiquitin (V Kirkin et al., 2009; Walinda et al., 2014). In the context of the recombinant protein, this deletion also abolished ubiquitin binding entirely (Fig 1G). When tested in the phase separation assay, the NBR1 PB1 deletion mutant displayed a severely reduced phase separation promoting activity (Figure 1H). The PB1 domain alone did not promote phase separation and in fact showed a dominant negative effect on the reaction (Fig 1I), consistent with its p62 filament-capping and shortening activity (Jakobi et al., 2020). This suggests that the interaction of the NBR1 PB1 domain with p62 is not sufficient for the promotion of phase separation but that this interaction mediates the recruitment of another biochemical activity of NBR1 to the p62 filaments. Consistently, we observed that the UBA domain deletion mutant of NBR1 also showed a severely reduced promoting activity in our assay (Fig 1H). Both the PB1 and the UBA deletion mutants were still recruited to the condensates (Fig 1H).

We therefore conclude that NBR1 aids in efficient cargo clustering of p62, likely by bringing its high affinity UBA domain (Walinda et al., 2014) to the p62 filaments via its PB1 domain.

### Live imaging of endogenously tagged NBR1 and p62 shows partial overlap of the two proteins

In order to compare our *in vitro* results regarding the promotion of p62-positive condensate formation by NBR1 to the naturally occurring process in cells, we tagged endogenous NBR1 at the N-terminus with an mScarlet-AID (auxin inducible degron) tag in HAP1 cells expressing endogenously tagged GFP-p62 (Fig 2A, B and Fig S2A, B). The fluorophores on p62 and NBR1 allowed us, for the first time, to image cargo-receptor interplay and dynamics at endogenous expression levels in live cells. mScarlet retains some stability under acidic conditions and we observed a band positive for mScarlet at approx. 30kDa which we interpret as the cleaved form of the fluorophore after NBR1 has been degraded within the lysosome (Fig 2B and Fig S2B). Treatment with Bafilomycin A, which blocks the acidification of the lysosome, led to stabilization of full length mScarlet-AID-NBR1 and a less prominent free mScarlet band (Fig 2B and Figure S2B).

**Figure 2.**
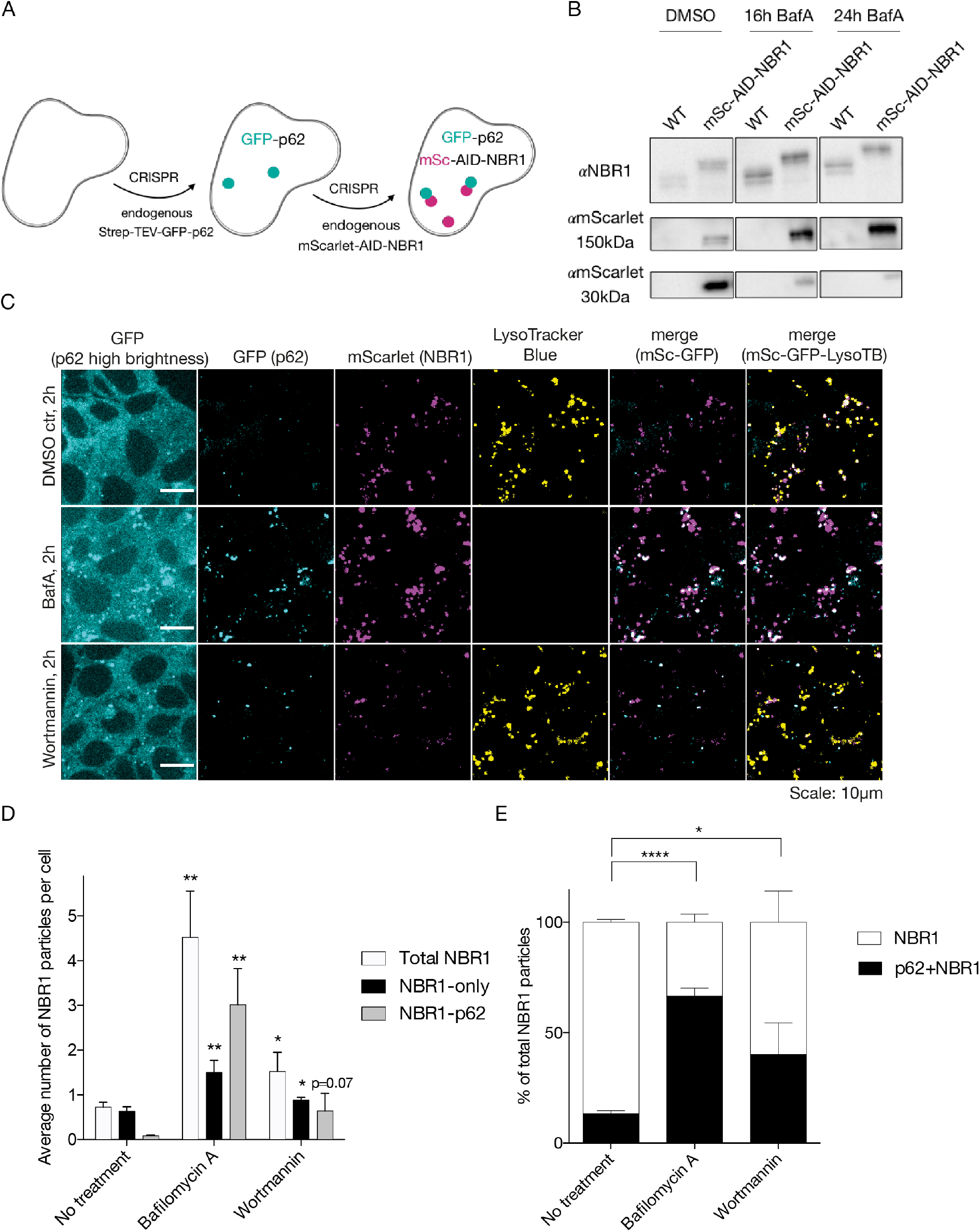
Distribution of endogenous mScarlet-tagged NBR1 in the cell. **(A)** Schematic of the generation of the used cell lines. Two subsequent CRISPR reactions introduce a Strep-TEV-GFP tag on endogenous p62 and an mScarlet-AID-tag on endogenous NBR1. **(B)** Western blot of 20 μg total protein lysate from endogenously N-terminally tagged NBR1 in HAP1 cells with mScarlet-AID. The upper panel shows an anti-NBR1 staining and the lower an anti-mScarlet staining for full length mScarlet-AID-NBR1 (140kDa) and likely free mScarlet (30kDa). Treatment with BafA was performed on a subset of the samples as indicated. The uncut membranes, containing several additional controls are shown in Fig S2A, B. **(C)** Live cell imaging of endogenously tagged mScarlet-AID-NBR1 cells. Note: mScarlet is acid-stable and emits a signal of potentially degraded mSc-AID-NBR1 protein from inside lysosomes. Note: for BafA treated cells the lysosome is de-acidified and the signal of p62 and NBR1 from the lysosome is unquenched. Therefore, these samples include GFP and mScarlet signals present in lysosomes. **(D)** Quantification of three replicates of imaging as shown in C. Quantification was done on one plane from an acquired Z stack by thresholding and counting the number of particles which partly or completely overlap. Plotted are the total number of NBR1 particles which do not overlap with LysoTrackerBlue signal and the subsets of total NBR1 particles which either colocalizes or not with GFP-p62. Significance (unpaired Student’s t-test) was calculated based on comparison with untreated samples for each type of particles. The average number of p62 particles for the same replicates is shown in Fig S2C. **(E)** Percentage of mSc-AID-NBR1 particles outside of the lysosomes (not colocalizing with LysoTracker Blue) which either colocalize with GFP-p62 or not.

Live imaging of the distribution of GFP-p62 and mScarlet-AID-NBR1 was complicated by the pH resistant mScarlet signal emitted from the lysosomes. We therefore excluded the mScarlet signal which overlapped with acidified vesicles, stained by LysoTracker Blue and considered only the signal which was outside of these compartments (Fig 2C). We quantified the total number of mScarlet-NBR1 particles outside of lysosomes and the number of these particles which colocalized with p62 (Fig 2C, D). A large population of NBR1 puncta did not overlap with p62 (Fig 2D, E). Upon treatment with wortmannin, which blocks autophagosome formation we observed an increase of p62 puncta (Fig S2C) and also a higher degree of colocalization of the two proteins (Fig 2E), suggesting that the double positive condensates are specifically turned over by autophagy. Consistently, treatment with bafilomycin A showed an increase in all populations of NBR1, most notably the population of NBR1 colocalizing with p62 (Fig 2D, E). However, these data should be evaluated carefully, since the treatment with bafilomycin A leads to an unquenching of both mScarlet and GFP signals present in the lysosome and simultaneously interferes with LysoTrackerBlue staining.

### Acute depletion leads to a rapid reduction of the number of p62 condensates in cells

Having established the cell line and conditions for the live imaging of NBR1 and p62, we stably introduced TIR1, an E3 ligase which upon stimulation with 1-NAA (1-naphtaleneacetic acid) is able to recognize the AID tag fused to NBR1 and induce its proteasomal degradation (Fig 3A, (K. Nishimura, Fukagawa, Takisawa, Kakimoto, & Kanemaki, 2009)). The NBR1 protein in the parental cell line in which no TIR1 was introduced was not affected by treatment with 1-NAA (Fig 3B, C). The TIR1-containing cell line, however, was able to deplete the majority of the NBR1 protein efficiently within 3 hours of treatment. (Fig 3B, C). Simultaneous treatment with 1-NAA and the proteasomal inhibitor MG132 led to stabilization of the protein suggesting that, as expected (K. Nishimura et al., 2009), TIR1/1-NAA-induced degradation of NBR1 is mediated by the proteasome. Despite being a prominent interactor of NBR1 *in vivo*, p62 was not collaterally degraded by the utilized degron system upon addition of 1-NAA (Fig 3B).

**Figure 3.**
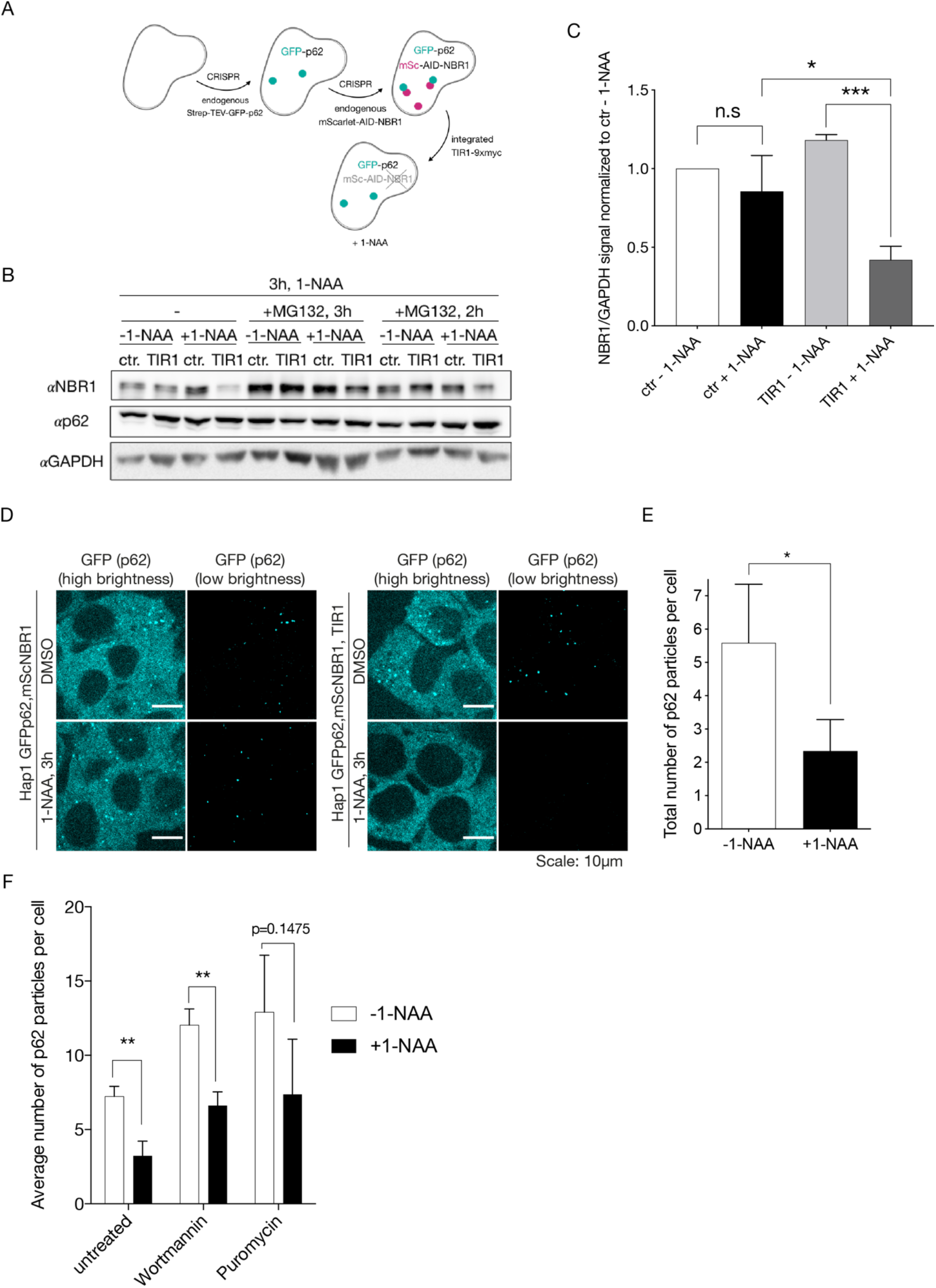
Acute NBR1 depletion leads to formation of less p62 puncta in cells. **(A)** Schematic of the generation of the used cell line. Stable introduction of TIR1 in the Strep-TEV-GFP endogenous tag on p62 and mScarlet-AID-tag on endogenous NBR1 allows subsequent depletion of endogenous NBR1. **(B)** Western blot and quantification showing depletion of endogenous mSc-AID-NBR1 upon treatment with 1-NAA. The control cell line is HAP1 GFP-p62, mScarlet-AID-NBR1, whereas the TIR1 cell line stably expresses the plant TIR1 ligase in addition. The cells were treated for a total of 3h with 1-NAA, parts of the samples were treated either simultaneously or for the final 2h prior to lysis with MG132. **(C)** Quantification of the western blot in B. **(D)** HAP1 GFP-p62, mScarlet-NBR1 cells without and with integrated TIR1-9xmyc were treated for 3h with 1-NAA or vehicle (DMSO). **(E)** Quantification of 3 experiments of NBR1 depletion. The average number of p62 puncta was quantified and plotted for cells containing TIR1. **(F)** GFP-p62, mScarlet-NBR1, TIR1 cells were treated with 1-NAA for a total of 3h and with wortmannin or puromycin for a total of 2h. The average number of p62 puncta was quantified. Quantification with standard deviation and statistics based on an unpaired Student’s t-test.

Next, we examined the effects of acute depletion of NBR1 on the number of p62 condensates as assessed by p62 positive puncta. We treated the cell line containing GFP-p62, mScarlet-AID-NBR1 and TIR1 (Fig 3A) with 1-NAA for 3 hours and subjected the cells to live cell imaging (Fig 3D). We observed that the number of GFP-p62 puncta was significantly reduced (Fig 3D, E). Treatment with wortmannin and puromycin promoted higher numbers of total p62 puncta, however, still showed a lower amount of p62 puncta compared to when the 1-NAA treatment was omitted (Fig 3F).

In conclusion, a rapid, targeted depletion of endogenous NBR1 from cells leads to a significantly reduced number of p62 puncta, indicating a potentially diminished capacity to cluster cargo for selective autophagy.

### Expression of PB1 and UBA domain mutants of NBR1 fail to rescue acute depletion of endogenous NBR1

Having established that acute depletion of endogenous NBR1 results in reduced condensate formation of endogenous p62 in living cells, we asked if the expression of NBR1 or its PB1 and UBA domain mutants could rescue the depletion of NBR1. To this end, we generated stable cell lines expressing doxycycline-inducible 3xflag-iRFP-NBR1 in the GFP-p62, mScarlet-AID-NBR1, stable-TIR1 background (Fig 4A). The NBR1 variants integrated into the cells were either wild type NBR1, a D50R mutant which shows reduced binding to p62 (V Kirkin et al., 2009; Lamark et al., 2003) or an F929A mutant which is defective in ubiquitin binding (Walinda et al., 2014). Stable clones were selected based on similar expression levels of iRFP-NBR1 variants after doxycycline treatment (Fig 4B). The efficiency of degradation of endogenous NBR1 in the selected clones upon 1-NAA addition showed some variability which was adjusted by treating with different 1-NAA concentrations, as indicated in the figure legend (Fig 4B). Despite of not observing any changes of the p62 levels in the cell lines between the clones on a western blot level (Fig 4B), the baseline levels of p62 puncta between the clones differed. In particular, the F929A cell line presented more p62 puncta than the wild type or D50R clones (Fig 4C, D and Fig 5). Taking this into consideration, we compared the number of p62 puncta and their area within each generated cell line at resting state, NBR1-depletion and doxycycline re-expression. Overexpression of wild type NBR1 rescued depletion of endogenous NBR1 by 1-NAA as we observed no significant difference between untreated cells and cells rescued with wild-type NBR1 in terms of p62 puncta number (Fig 4C, D). The area of the puncta, however, appeared to be larger when NBR1 was overexpressed (Fig 4C, E), consistent with a previous report (Sánchez-Martín et al., 2020).

**Figure 4.**
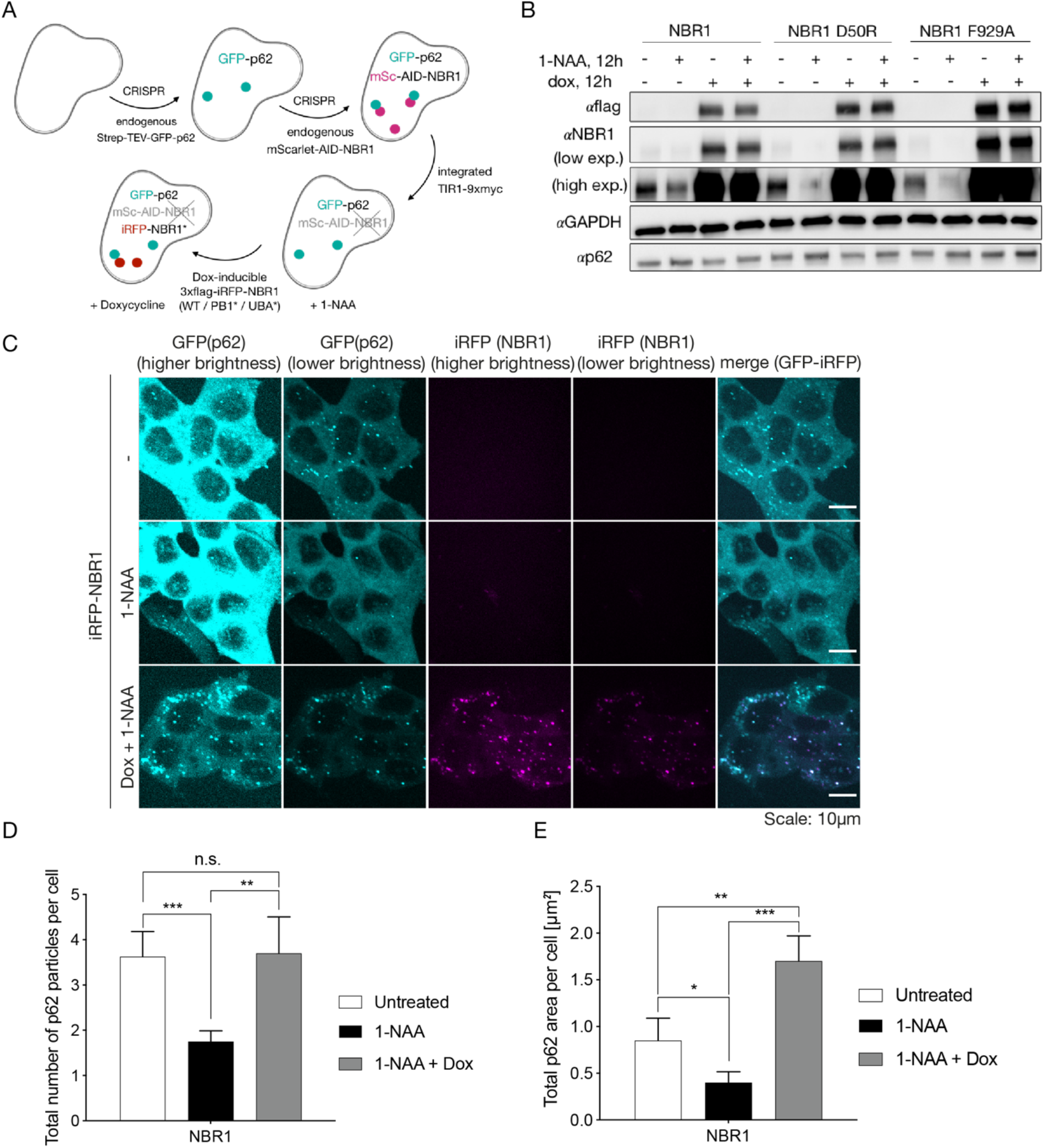
Re-introduction of NBR1 carrying PB1 or UBA inactivating mutations after depletion of endogenous NBR1 does not rescue p62 puncta formation. **(A)** Schematic of the generation of the used cell lines. Stable introduction of iRFP-NBR1 variants under a doxycycline-inducible promoter allows rescue experiments while the endogenous NBR1 is depleted and the endogenous p62 can be imaged via the integrated GFP fluorophore. **(B)** Western blot showing endogenous NBR1 depletion for the generated cell lines upon 1-NAA treatment (1mM for NBR1 and 500µM for the NBR1 D50R and F929A mutant cell lines), re-expression of 3xflag-iRFP-NBR1 variants upon treatment with 50ng/ml doxycycline and combined treatment. **(C)** Representative images of the experiment depicting p62 puncta from a Z-stack maximum signal projection in the WT NBR1 rescue cell line when treated with 1-NAA or 1-NAA and doxycycline. **(D, E)** Quantification of four replicates depicting the total number **(D)** or total area **(E)** of p62 puncta from the images shown in 4C.

**Figure 5.**
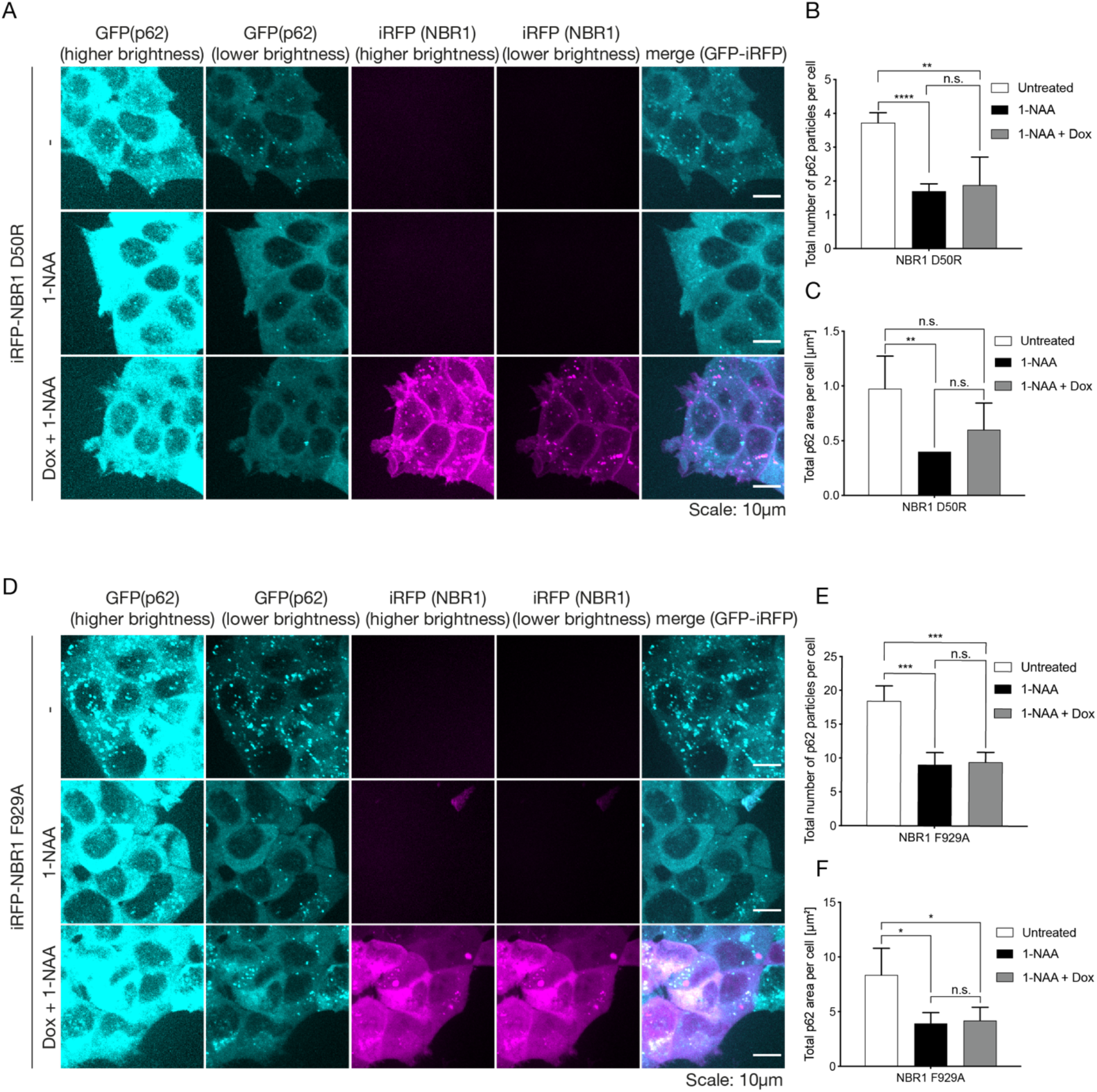
Re-introduction of NBR1 carrying PB1 or UBA inactivating mutations after depletion of endogenous NBR1 does not rescue p62 puncta formation. **(A)** Representative images showing p62 puncta from a Z-stack maximum signal projection in the NBR1 D50R rescue cell line when treated with 1-NAA or 1-NAA and doxycycline. **(B, C)** Quantification of four replicates depicting the total number **(B)** or total area **(C)** of p62 puncta from the images shown in 5A. **(D)** Representative images from the same experiment performed for the NBR1 F929A rescue cell line as in 5A and B. **(E, F)** Quantification of four replicates of the images shown in 5D for the number of p62 puncta **(E)** or area **(F)**.

Performing the same treatments for the PB1-mutated D50R cell line we observed a different pattern of NBR1 re-expression (Fig 5A). The signal for iRFP-NBR1D50R was much more diffuse and the formed puncta were smaller and less prominent. Quantification showed that in contrast to the wild type protein (Fig 4C, D) the overexpression of the D50R mutant of NBR1 was not able to rescue the number of p62-puncta compared to pre-depletion of endogenous NBR1 (Fig 5B).

The UBA-mutant F929A cell line, similarly to the PB1-mutant cell line, was not able to rescue the p62-puncta phenotype upon 1-NAA-mediated depletion of endogenous NBR1 and mutant re-expression. The overexpressed iRFP-NBR1 F929A protein showed a diffuse pattern of signal with very few large puncta (Fig 5D). The levels of p62 puncta were fewer than the pre-treatment levels and similar to when there was no doxycycline-mediated re-expression (Fig 5E).

We conclude that the PB1 and UBA domains of NBR1 are required to promote p62 condensate formation in cells and regulate their number and size.

## Discussion

Here we have shown, employing a fully reconstituted system and endogenously tagged proteins coupled with acute protein depletion, that the ability of NBR1 to promote phase separation of p62 and ubiquitinated cargo depends on its PB1 and UBA domains. p62 mediates the degradation of misfolded, ubiquitinated proteins within the lysosomes by selective autophagy (Bjørkøy et al., 2005; Danieli & Martens, 2018). This entails the phase separation of p62 filaments with ubiquitinated substates (Ciuffa et al., 2015; Jakobi et al., 2020; Sun et al., 2018; Zaffagnini et al., 2018), followed by the recruitment of the autophagy machinery (Turco et al., 2019) and finally attachment of the cargo to the nascent autophagosomal membrane (Ichimura et al., 2008; Pankiv et al., 2007; Wurzer et al., 2015). During these processes, and in particular during cargo condensation, p62 is assisted by NBR1 (V Kirkin et al., 2009; Sánchez-Martín et al., 2020; Zaffagnini et al., 2018) but the precise mechanisms of NBR1’s contribution are unclear. The PB1 and UBA domains of NBR1 have previously been implicated in the regulation of p62-mediated selective autophagy of ubiquitinated proteins (Jakobi et al., 2020; V Kirkin et al., 2009; Sánchez-Martín et al., 2020). However, if they directly impact the ability of p62 to phase separate with ubiquitinated cargo was unknown. The results of our reconstituted system suggest that the PB1 mediated recruitment of NBR1 to p62 is required to promote condensate formation. The PB1 domain of NBR1 itself is not sufficient to promote condensate formation and even inhibits the reaction when added on top of wild type NBR1. This suggest that NBR1 harbours other biochemical properties which upon recruitment to p62 promote phase separation. Our results further show that the NBR1 UBA domain confers one of these activities as the UBA deletion mutant showed a severely reduced promoting effect. It appears that the UBA domain mutant must be linked to the p62 filaments in order to promote phase separation because the PB1 mutant of NBR1 is still recruited to the condensates, possibly via the UBA domain mediated interaction with ubiquitin but nevertheless shows reduced facilitation of phase separation. The UBA domain of NBR1 has a higher affinity for ubiquitin compared to the UBA of p62 (Long et al., 2010; Walinda et al., 2014) and thus the heterooligomeric NBR1-p62 complex may have a higher affinity for ubiquitinated substrates than homooligomeric p62 filaments and therefore induce phase separation at lower substrate concentrations. Interestingly, at least *in vitro*, the phase separation promoting activity of the NBR1 was not completely lost upon deletion of the NBR1 UBA domain. NBR1 itself dimerizes or oligomerizes (Pankiv et al., 2007; Zaffagnini et al., 2018). It is therefore conceivable that NBR1 caps and shortens p62 filaments via its PB1 domain (Jakobi et al., 2020) but that the heterooligomers have a non-filamentous shape that is more efficient in phase separation.

A further observation we made was that a population of the endogenous NBR1 appears to be distributed in the cell without colocalizing with p62 puncta. This may hint at p62-independent NBR1 functions. It was proposed that NBR1 participates in a form of microautophagy in which the cargo is directly engulfed by the vacuole, irrespective of the rest of the autophagy machinery (Liu et al., 2015). NBR1 has also been shown to function in pexophagy (Deosaran et al., 2013). Endosomal microautophagic clearance of cargo receptors, including NBR1, has been reported under conditions of starvation (Mejlvang et al., 2018).

The question of cooperation of p62 and NBR1 in various autophagic processes needs further examination, as it is unclear whether NBR1 can contribute to downstream steps of the autophagic process with its additional domains which include two highly charged coil-coil domains, a conserved four tryptophan domain and a Jn domain, suggested to bind to membranes (Mardakheh, Auciello, Dafforn, Rappoport, & Heath, 2010).

## Acknowledgements

We thank Eleonora Turco and Luca Ferrari for comments on this manuscript. This work has been funded by the Austrian Science Fund (FWF P30401-B21 and F79) and a Uni:docs fellowship of the University of Vienna.

## Competing interest

Sascha Martens is member of the scientific advisory board of Casma Therapeutics.

## Materials and methods

### Plasmids and cloning

The plasmids pETDuet-mCherry-p62, pGEX-GST-4xUb and pFastBacHTB-GFP-NBR1 were previously described in Zaffagnini et al., 2018.

Cloning of gRNAs for CRISPR in the all-in-one plasmid containing Cas9 was done via BbsI for guide 1 and subsequently with BsaI for guide 2. The NBR1 homology template was generated from isolated gDNA (GeneJET Genomic DNA Purification Kit, Thermo Scientific, cat# K0721) and cloned into a pUC19 vector. The mScarlet and AID tags were amplified with homology primers for Gibson cloning and inserted into the pUC19-NBR1 homology template plasmid.

Cloning of NBR1 variants in pInducer20 was done in several steps. iRFP was cloned into a p3xflag-containing vector with KpnI and NotI. NBR1 variants (WT, PB1D50R and UBAF929A) carrying a short non-charged linker to the N-terminus were amplified and cloned after iRFP with KpnI and AgeI. Finally, the generated sequences 3xflag-iRFP-NBR1 variants were cloned via a Gibson cloning strategy into pInducer20.

All constructs were sequenced thoroughly before expression in bacteria or transfection into cells. All constructs used for mammalian cell transfection, including CRISPR-related plasmids, were purified using an endonuclease free plasmid extraction kit (NucleoBond Xtra Maxi Plus EF, Macherey-Nagel, cat# 740426.10).

### Protein expression and purification

mCherry-p62, GST-4xUb and GFP-NBR1 were expressed as previously described in Zaffagnini et al., 2018. GFP-NBR1ΔPB1 and GFP-NBR1ΔUBA were cloned and expressed in the same manner as WT GFP-NBR1.

*In vitro* synthesis of K48- and K63-linked ubiquitin chains was performed as previously described in Zaffagnini et al., 2018.

### Direct interaction assays

GST-4xUb or GST, as a negative control, proteins were recruited to GSH beads and the proteins tested for interaction (GFP-NBR1 variants) were added. Direct interaction is visualized by the appearance of a fluorescent signal on the rim of the beads. The proteins used to coat the beads, GST or GST-4xUb, were spun for 15 min at maximum speed to eliminate precipitates. 10 μl of GSH bead slurry (Pierce Glutathione Agarose Thermo Scientific, cat# 16101) was used per conjugated bead type. The beads were washed 3 times in SEC buffer (25mM Hepes pH 7.5, 150mM NaCl, 1mM DTT) and spun at 1000g for 1 min to remove the supernatant. A total of 25 μg protein for coating was added to the beads for 30 min/4^°^C/rolling overhead. The beads were washed in SEC buffer and added to 394-well imaging plates. The tested GFP-NBR1 variants were also spun after thawing to remove aggregates and a final concentration of 2 μM was added to the tested wells. The beads were imaged at a confocal Zeiss LSM700 microscope. After imaging, the samples were taken up and mixed with an SDS-loading dye and boiled for 5 min at 95 ^°^C. They were then loaded on a 10% SDS-PAGE gel and stained with Coomassie to control for equal input.

For the analysis, an average of 50 beads or more were imaged in Z stack of 45 μm. A maximum projection of the signal was made in Fiji and 6 lines were drawn per bead to quantify the maximum signal on the bead surface. The signal of 50 beads per sample was averaged. 3 replicates were performed for each condition and the averages from these replicates was given as input in Prism and ultimately plotted as graphs.

Testing of the interaction of the NBR1 variants with p62 was performed in a similar manner by conjugating a total of 25ug mCherry-p62 to RFP trap beads (Chromotek, rta-20).

### Phase separation assays

The assay was performed as previously described in Zaffagnini et al., 2018. In brief, 2 μM mCherry-p62 was premixed with 2 μM GFP-NBR1 in SEC buffer (25mM Hepes pH 7.5, 150mM NaCl, 1mM DTT). The reaction was triggered upon addition of simulated cargo, 5 μM GST4xUb or K48/K63-linked chains. Alterations to the buffer conditions and concentrations of NBR1 or additional components are specified in the figure legends. Due to variability of the baseline of the control reaction containing p62 and GST4xUb in terms of the peak particle number, in some experiments a crowding agent such as 2% BSA was introduced into the buffer before the addition of ubiquitin or ubiquitinated cargo. The presence of up to 5% BSA in the buffer did not trigger condensate formation in the absence of simulated cargo containing ubiquitin. Time-lapse imaging was started when the cargo was added and imaging was performed every minute for 1 hour. Quantification of condensate number and size were made based on mCherry fluorescence unless otherwise specified.

### Tissue culture

HAP1 cells from Horizon Discovery were grown in Iscove’s Modified Dulbecco’s Medium (Gibco GlutaMAX, Thermo Fisher Scientific) with 10% fetal bovine serum (Sigma Aldrich, F7524) and Penicillin-Streptomycin (10 000U/ml Penicillin and 10mg/ml Streptomycin, Sigma Aldrich, P4333). The cells were split every 2 days.

### CRISPR

To endogenously tag NBR1 with an mScarlet-AID or GFP-AID tags, the genomic area approx. 1500bp up and downstream of the ATG start codon was amplified from isolated HAP1 genomic DNA (obtained using GeneJET Genomic DNA Purification Kit, cat#K0721, Thermo Fisher Scientific) and sub-cloned into pUC19 (Addgene, #50005). The corresponding tags, mScarlet, GFP and AID were amplified via PCR and assembled via Gibson cloning (using the Gibson Assembly Master Mix, cat# E2611, NEB). The resulting plasmid was sequenced thoroughly. The chosen gRNAs were cloned into an all-in-one (AIO) plasmid containing Cas9D10A nickase via standard BbsI cleavage sites and the resulting plasmid was sequenced. The AIO plasmid and the template plasmid were transfected into HAP1 STG-p62 (for mScarlet-AID-NBR1) or HAP1 WT (for GFP-AID-NBR1) cells using FuGene 6 transfection reagent (Promega, cat# e2691). 48h later the cells were sorted for mScarlet fluorescence in bulk, left to expand until confluent in a 15cm dish (approx. 10 days) and then resorted for mScarlet fluorescence as single clones. The sorted single clones were left to expand in 96-well plates for approx. 10 days and split to 3 separate 96-well plates. One plate was frozen, one plate left to expand and one plate was used for a genotyping PCR (primers annealing in the homology template around the ATG. If negative, a band of 350bp would have been amplified and if the tags were present, a band of 1kb would have been amplified. The PCR was performed with REDTaq ReadyMix (Sigma-Aldrich, cat# R2523) and loaded on a 1% agarose gel. At least 6 positive clones per cell line were further expanded and tested on a gDNA, cDNA and western blot levels. gDNA was isolated and the area containing the tag was amplified with primers outside of the homology area amplified for the CRISPR reaction. The resulting PCR product was sequenced. cDNA was generated from the gDNA (via First Strand cDNA Synthesis Kit, Roche, cat# 11483188001) and a PCR was done with primers starting at mScarlet and the end of NBR1. The resulting PCR was sub-cloned into pETDuet and thoroughly sequenced. Cell lysates from the clones were loaded on SDS-PAGE for western blotting and probed for NBR1, mScarlet and Cas9. All the clones showed a shift due to the introduction of the tag. None of the clones had integrated Cas9. One clone was chosen to proceed with further experiments.

### Stable cell line generation

A stable cell line which depletes endogenous mScarlet-AID-NBR1 was generated by introducing a TIR1-9xmyc lentiviral plasmid which integrated randomly in the genome. The cells were transfected using a standard protocol with FuGene 6 transfection reagent. They were selected with puromycin (5 μg/ml) until the control untransfected cells died. The cells were then single sorted to 96-well plates and left to expand. Initial screening was done on a PCR level by amplifying a short fragment of the inserted proteins. Validation was done on a western blot level for the myc tag and for NBR1 depletion upon treatment with 1-NAA.

The stable cell lines expressing 3xflag-iRFP-NBR1 variants were generated in a similar fashion by selection with G-418 (1200 μg/ml). They were sorted based on iRFP fluorescence after a 24h treatment with 500ng/ml doxycycline. After PCR screening, they were tested with various doxycycline concentrations for various periods of time to ensure similar expression levels of the flag-tag and NBR1 levels.

### Cell treatments, western blot and preparation for live cell imaging

Cells plated for live cell imaging (5 000 cells were plated in imaging chambers (Greiner Bio One, cat# 543079)) or western blot analysis (200 000 cells in 6-well plates (Eppendorf, cat# 0030720113) were left to attach for 48h. Treatments with 500 μM 1-NAA were done for 3 hours (tested 1-, 4- and 24-hour time points). If other treatments were performed on top, they were done for 2 hours within this period of 3-hour 1-NAA treatment. Used concentrations for the treatments are as follows: puromycin 0.5 μg/ml, MG132 10 μM, wortmannin 1 μM, bafilomycin A 0,4 μM.

For western blot: At the end of the 3-hour incubation period cells were washed 3x PBS and lysed for western blot in lysis buffer (20 mM Tris pH 8, 10% glycerol, 135 mM NaCl, 0.5% NP40, 2.5 mM MgCl2, DNAse (Sigma-Aldrich, cat# DN25-1G), 1x protease inhibitors (cOmplete Protease Inhibitor Cocktail, Roche, cat# 11697498001)). The cells were scraped from the plate in lysis buffer, left on ice for 10 min, spun down at maximum speed in a tabletop centrifuge for 10 min and the supernatant was transferred to a fresh tube. Equal loading was ensured by conducting protein concentration measurements with the Pierce BCA Protein Assay Kit (Thermo Scientific, cat# 23225). After SDS-PAGE and western blot transfer, the membranes were stained with Ponceau S (Sigma-Aldrich, cat# P3504) and blocked with 3% milk in TBST buffer for 30 min at room temperature. The membranes were then incubated with primary antibodies overnight at 4^°^C. After 3x 5 min washes in TBST, secondary antibodies were added for 30 min at room temperature and the membranes were washed 3x 5 min in TBST again and developed with ECL (SuperSignal™ West Pico PLUS Chemiluminescent Substrate, Thermo Scientific, cat#34580 or SuperSignal™ West Femto Maximum Sensitivity Substrate, Thermo Scientific, cat# 34095).

For live cell imaging: 30 min prior to the end of the 3-hour incubation time point the media was exchanged to media with the corresponding treatments and 0.1 μM LysoTrackerBlue. At the end of the time point the media was exchanged to media with the corresponding treatments without LysoTrackerBlue and the cells were imaged.

### Live cell imaging and analysis

Cells were imaged in a temperature- and CO_2_-controlled environment with a Visitron Live Spinning Disk (Plan-Apochromat 63x/1.4 Oil DIC objective and an EM-CCD camera). As standard imaging conditions GFP was imaged with 10% laser power, mScarlet with 14%, LysoTrackerBlue with 3% and 10% for iRFP. Imaging conditions were kept consistent within the same experiment and among replicates. A stack of 5 μm was taken for each image with a 0.5 μm distance. 11 slices were taken per condition.

The analysis for colocalization was performed on a single plane to exclude structures positioned on top of each other instead of on the same plane. For total particle numbers per cell, a maximum signal projection was quantified. The colocalization and quantification were performed by setting an arbitrary threshold for each channel. The resulting thresholded image was visually assessed for correct representation of puncta in the cells as compared to the original image. The thresholds were kept consistent within the same experiment, slight adjustments were made among replicates due to potential variations of laser strength at different days when the experiments were repeated.

For total number of particles: A macro was designed to quantify the number of particles and total particle area for each channel by using the “Analyze particles” function in Fiji. The number of cells per image were manually counted and the particle number was divided by the cell number to get the average number of particles per cell. This was done for a maximum projection of all particles within the acquired 11-image stack.

For colocalization: A macro was designed to quantify the number of particles and total particle area for each channel. The selected ROIs from this macro were then overlapped with the thresholded image from another channel and depending on the end-goal of the analysis, if the ROIs detected signal or did not detect a signal, they were counted further for overlap in another channel. In this manner, mScarlet particles (representing NBR1) which did not colocalize with lysosomes (LysoTrackerBlue) could be counted as either colocalizing with GFP (p62 particles) or not.

The values were entered into Excel and the average for 3 replicates was calculated. The averages were copy-pasted in Prism (GraphPad Prism 8) and plotted as graphs. An unpaired Student’s t-test was performed to assess statistical significance. P values are presented with asterisks in the following cases: P<0.05 *, P<0.01 **, P<0.001 ***, P<0.0001 **** The figure legends indicate among which samples the significance was calculated.

### List of used reagents

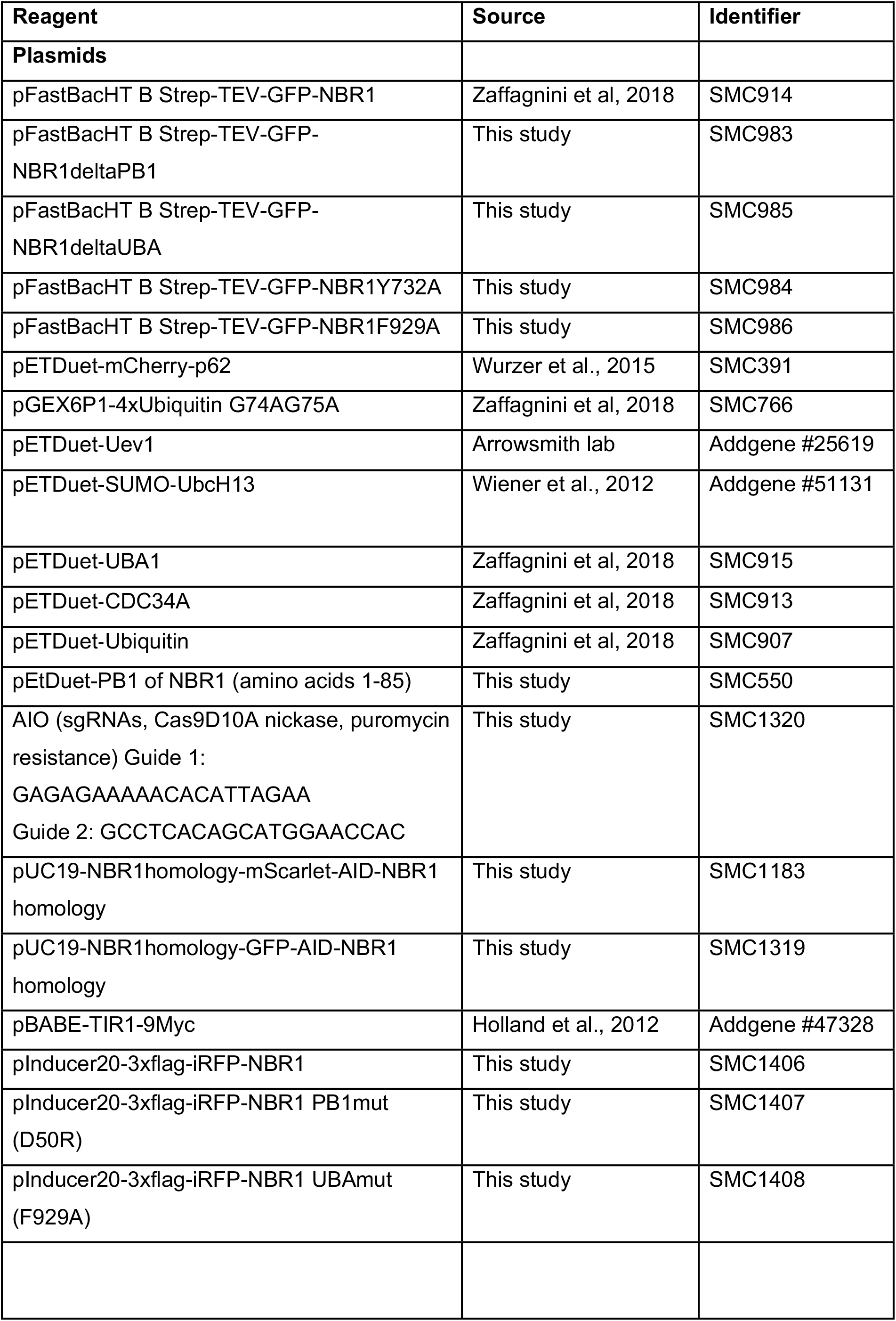
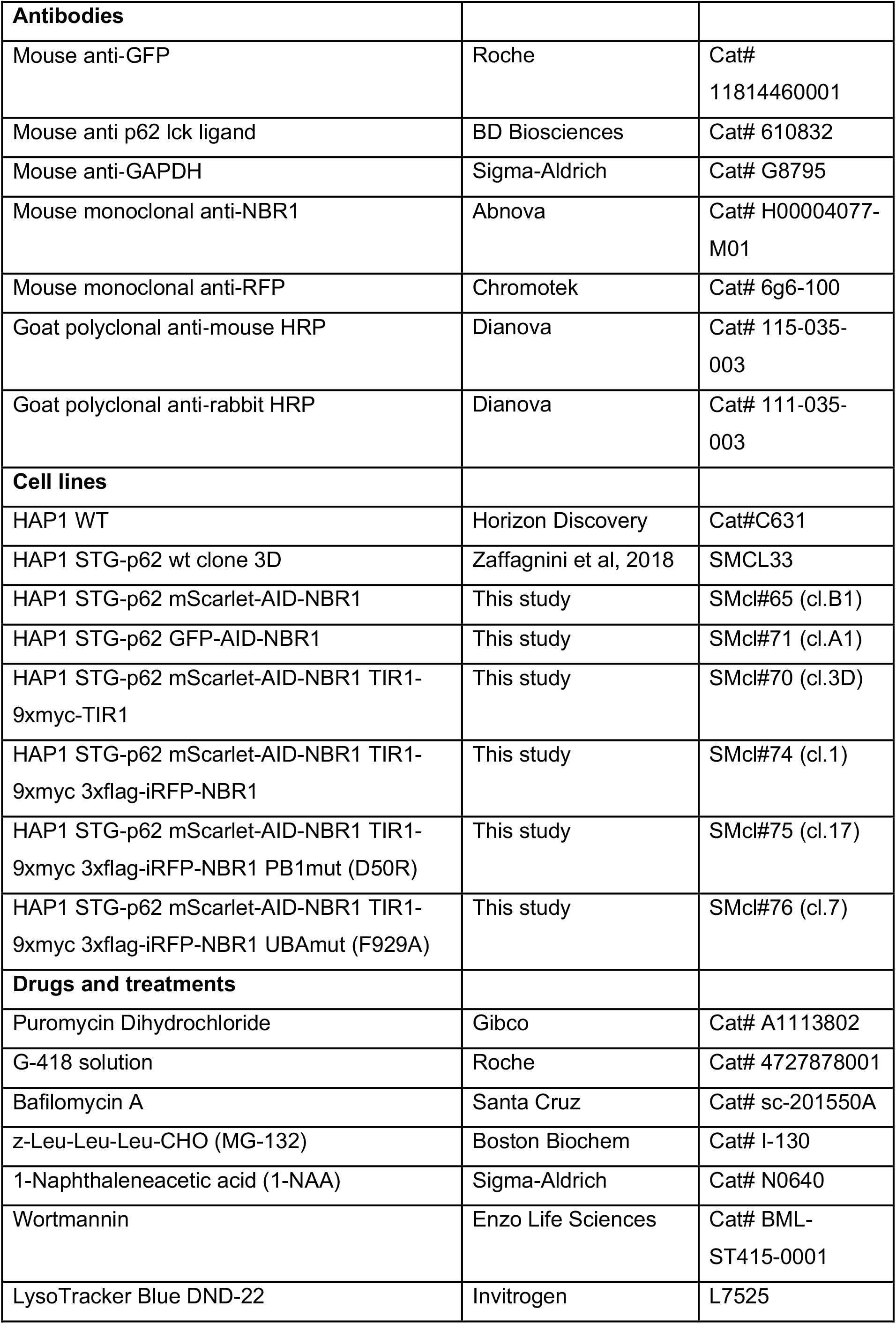

---

**Figure S1 – Figure supplement 1.**
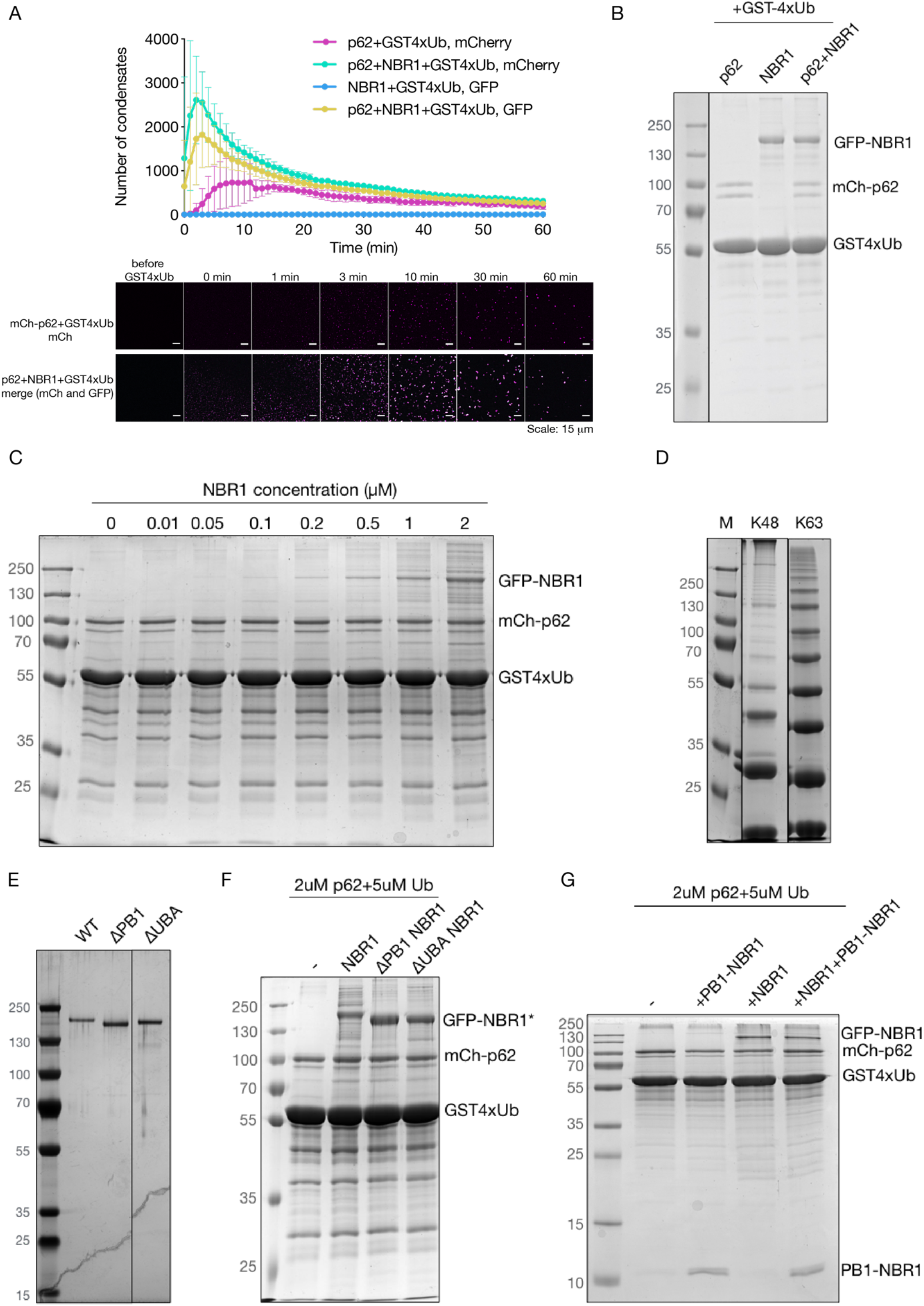
**(S1A related to Fig 1C)** The graph from Fig1C is shown with included quantification based on the GFP signal. Representative images from selected timepoints are displayed for the mCh-p62 + GST4xUb reaction (in mCherry) and mCh-p62 + GFP-NBR1 + GST4xUb reaction (merge for mCherry and GFP). **(S1B related to Fig 1C)** The volume of the phase separation reaction from Fig1C was loaded on a 10% SDS-PAGE and stained with Coomassie Blue. The middle reaction was not plotted in 1C, instead it is plotted in S1A, since the reaction GFP-NBR1+GST4xUb does not contain an mCherry protein and can only be quantified based on the GFP signal. **(S1C)** The volume of the phase separation reaction from Fig1D was loaded on a 10% SDS-PAGE and stained with Coomassie Blue. **(S1D)** Coomassie gel of the in-house synthesized K48- and K63-linked ubiquitin chains. **(S1E)** Silver staining of a polyacrylamide gel showing 1ug of purified recombinant GFP-NBR1 variants. Full length protein (WT) as well as PB1 domain and UBA deletion mutants are loaded. **(S1F, G)** The volume of the phase separation reaction from Fig 1H and 1I were loaded on a 10% SDS-PAGE and stained with Coomassie Blue.

**Figure S2 – Figure supplement 2.**
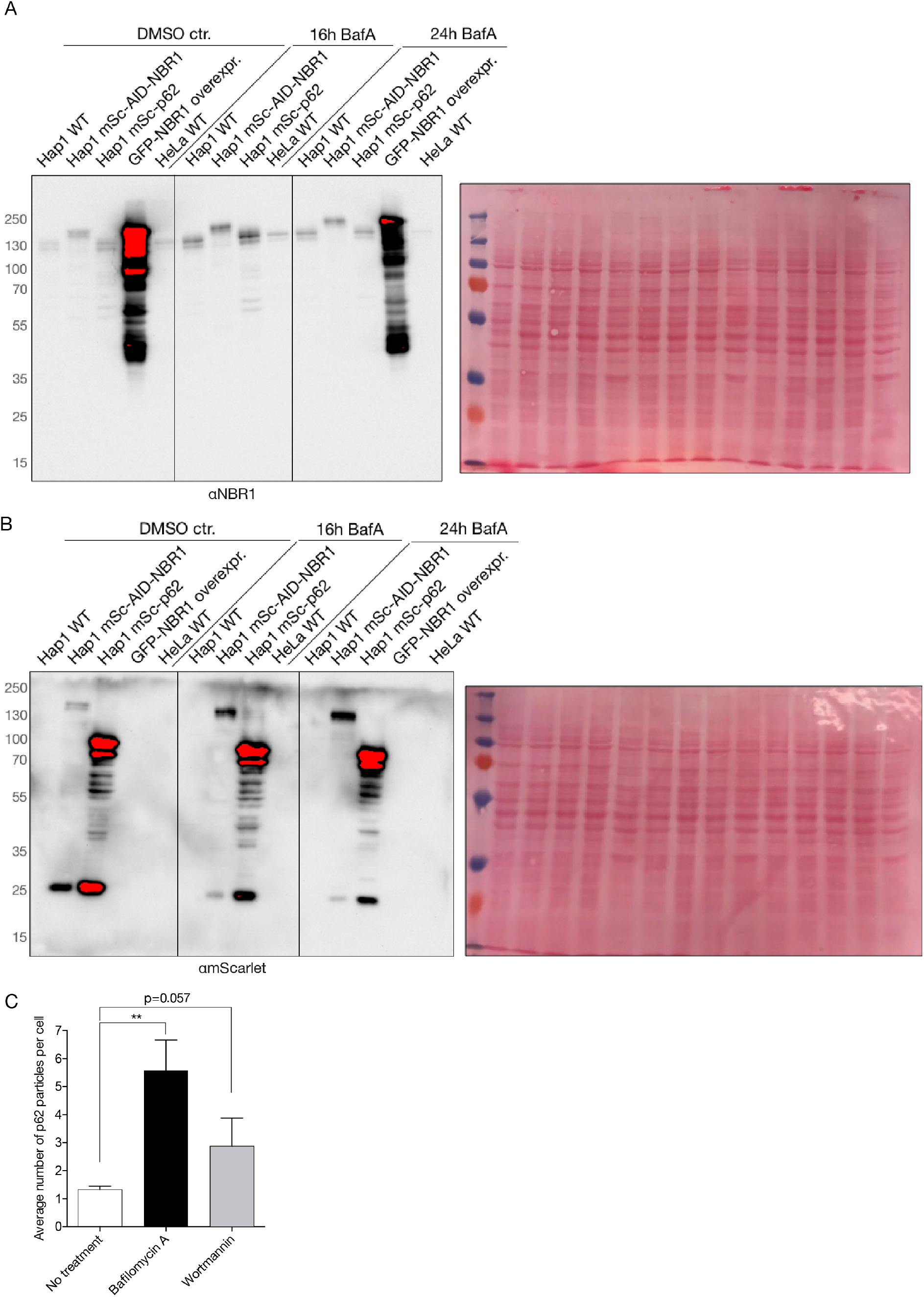
**(S2A, S2B for 2B)** Uncut western blots and Ponceau S staining for NBR1 and mScarlet as shown in 2B. Several controls are loaded, such as HAP1 mScarlet-p62 cell lysate, a lysate from a transiently transfected HAP1 WT cell with GFP-NBR1 and HeLa WT cells. **(S2C)** Quantification of total p62 particles from one plane of a Z-stack under treatments with BafilomycinA and Wortmannin from three replicates of imaging as shown in 2C.

